# Cell type-specific impact of aging and Alzheimer disease on hippocampal CA1 perforant path input

**DOI:** 10.1101/2024.08.27.609952

**Authors:** Bina Santoro, Kalyan V. Srinivas, Isabel Reyes, Chengju Tian, Arjun V. Masurkar

## Abstract

The perforant path (PP) carries direct inputs from entorhinal cortex to CA1 pyramidal neurons (PNs), with an impact dependent on PN position across transverse (CA1a–CA1c) and radial (superficial/deep) axes. It remains unclear how aging and Alzheimer disease (AD) affect PP input, despite its critical role in memory and early AD. Applying *ex vivo* recordings and two-photon microscopy in slices from mice up to 30 months old, we interrogated PP responses across PN subpopulations and compared them to Schaffer collateral and intrinsic excitability changes. We found that aging uniquely impacts PP excitatory responses, abolishing transverse and radial differences via a mechanism independent of presynaptic and membrane excitability change. This is amplified in aged 3xTg-AD mice, with further weakening of PP inputs to CA1a superficial PNs associated with distal dendritic spine loss. This demonstrates a unique feature of aging-related circuit dysfunction, with mechanistic implications related to memory impairment and synaptic vulnerability.

## INTRODUCTION

Memory decline is a hallmark symptom of normal aging and Alzheimer disease (AD), correlating with circuit dysfunction in the entorhinal-hippocampal network. Prior work on the circuit basis of aging- and AD-related memory loss has primarily highlighted the vulnerability of the trisynaptic pathway, the serial sequence of excitatory connections entorhinal cortex (EC) to dentate gyrus (DG) to CA3 and then to CA1 pyramidal neurons (EC→DG→CA3→CA1) (Small et al., 2011; Pozueta et al., 2013; Llorens-Martin et al., 2014). Yet, considerably less is known about how aging and AD impact the direct EC→CA1 inputs carried by the perforant path (PP), despite the critical role of this synapse in memory-guided behavior (Remondes and Schuman, 2004; Brun et al., 2008; Kitamura et al., 2014), and evidence of the sequential involvement or propagation of AD pathology from EC to CA1 early in the disease (Braak and Braak, 1991; Lace et al., 2009; de Calignon et al., 2012; Liu et al., 2012).

Moreover, to date, most aging and AD studies have analyzed CA1 pyramidal neurons (PN) as a homogeneous population, despite mounting evidence for the synaptic and intrinsic heterogeneity of CA1 PN circuits. Specifically, with regard to the PP, we previously demonstrated that the source and strength of direct EC inputs depend on the anatomical location of CA1 PNs across the transverse (CA2-subiculum) and radial (deep-superficial) axes (Masurkar et al., 2017). In CA1c, towards CA2, PNs are preferentially excited by medial EC (MEC) axons, with deep PNs as the primary target. In contrast, in CA1a, towards subiculum, PNs are preferentially excited by lateral EC (LEC) axons, with superficial PNs as the primary target. This input segregation supports *in vivo* evidence for the functional specialization of CA1c and deep PNs (dPNs) for spatial memory and CA1a and superficial PNs (sPNs) for non-spatial memory (Henriksen et al., 2010; Hartzell et al., 2013; Oliva et al., 2016; Burke et al., 2011; Ito and Schuman, 2012; Nakamura et al., 2013; Igarashi et al., 2014; Li et al., 2017), reflecting the respective roles of MEC and LEC in spatial and non-spatial memory (Fyhn et al., 2004; Hargreaves et al., 2005; Deshmukh and Knierim, 2011; Tsao et al., 2013; Igarashi et al., 2014). Additionally, we and others have also observed a heterogeneity of intrinsic excitability across these PN subpopulations that promotes this functional specialization (Maroso et al., 2016; Masurkar et al., 2020). Consistent with such specialization, PN subpopulations feature distinct gene expression profiles (Dong et al., 2009; Cembrowski et al., 2016) and associate with distinct interneuron subtypes (Lee et al., 2014; Valero et al., 2015).

This diversity may predict cell type-specific changes in response to aging and AD (Masurkar, 2018). Clarifying if and how these changes stratify across functionally specialized subpopulations would provide better cellular- and circuit-level insight for understanding *in vivo* findings. For example, *ex vivo* studies indicate that CA1 PNs undergo excitability changes with aging that are associated with cognitive deficits, but their correlation to *in vivo* population activity is inconsistent (Samson and Barnes, 2013). Additionally, age-related compromise of place cell remapping appears to be differentially expressed across the transverse CA1 axis (Hartzell et al., 2013), yet the synaptic basis is unclear.

Thus, several questions arise. Does aging differentially impact PP inputs across CA1 PN subpopulations? Are these changes mirrored in the terminal synaptic connections of the trisynaptic pathway, ie, the CA3→CA1 Schaffer collateral (SC) synapses? Does intrinsic excitability change in a way to amplify or compensate for these changes? And lastly, does AD pathology enhance this pattern? To address these questions, we applied a modified slice preparation technique that enabled robust *ex vivo* recordings from dorsal hippocampal slices from mice up to 30 months of age. We performed whole cell patch clamp recordings during extracellular stimulation of PP and SC inputs to CA1 PNs to interrogate excitatory input responses and intrinsic properties. We also applied 2-photon microscopy to quantify potential dendritic spine loss associated with these two synaptic pathways.

## RESULTS

### Aging has a distinct, cell type-specific impact on PP versus SC excitation

The CA1 PN apical dendrite receives input in compartmentalized fashion. PP inputs course through *stratum lacunosum moleculare* (SLM) and target the distal dendrites whereas SC inputs traverse *stratum radiatum* (SR) and synapse on the more proximal dendrites. In acute slices of mouse dorsal hippocampus (see Methods) derived from adult (3-6 month) and aged (24-30 month) C57Bl/6J mice, we performed whole cell patch clamp recordings from PNs subpopulations across the radial and transverse axes. We applied a modified slice preparation featuring intracardiac perfusion (see Methods), in both adult and aged mice, which enabled healthy recordings even in the oldest mice.

Recording from sPNs and dPNs in CA1c and CA1a, we examined PP-evoked synaptic responses using a range of stimulation intensities through an extracellular electrode in SLM to establish input-output curves (Figure 1Ai). Experiments were first performed with inhibition intact to examine the postsynaptic potential (PSP) that represented the summed response from the excitatory postsynaptic potential (EPSP) and the inhibitory postsynaptic potential (IPSP) (Figure 1Aii). In young mice (6-8 week), we previously found a striking difference in the relative strength of the PP PSPs in sPNs compared to dPNs that reversed along the transverse axis (Masurkar et al., 2017). In adult mice, as in young mice, the CA1c PP PSP was much larger in dPNs compared to sPNs (Figure 1B, p = 0.0002) whereas in CA1a the PP PSP was much greater in sPNs compared to dPNs (Figure 1C, p < 0.0001). Remarkably, aging abolished this differential response along the radial axis in both CA1c and CA1a by exerting opposing effects on the dPN and sPN PSPs (Figure 1B, C). Thus, in CA1c aging reduced the PP PSP in dPNs while it increased the PP PSP in sPNs, with the converse effect seen in CA1a.

**Figure 1.**
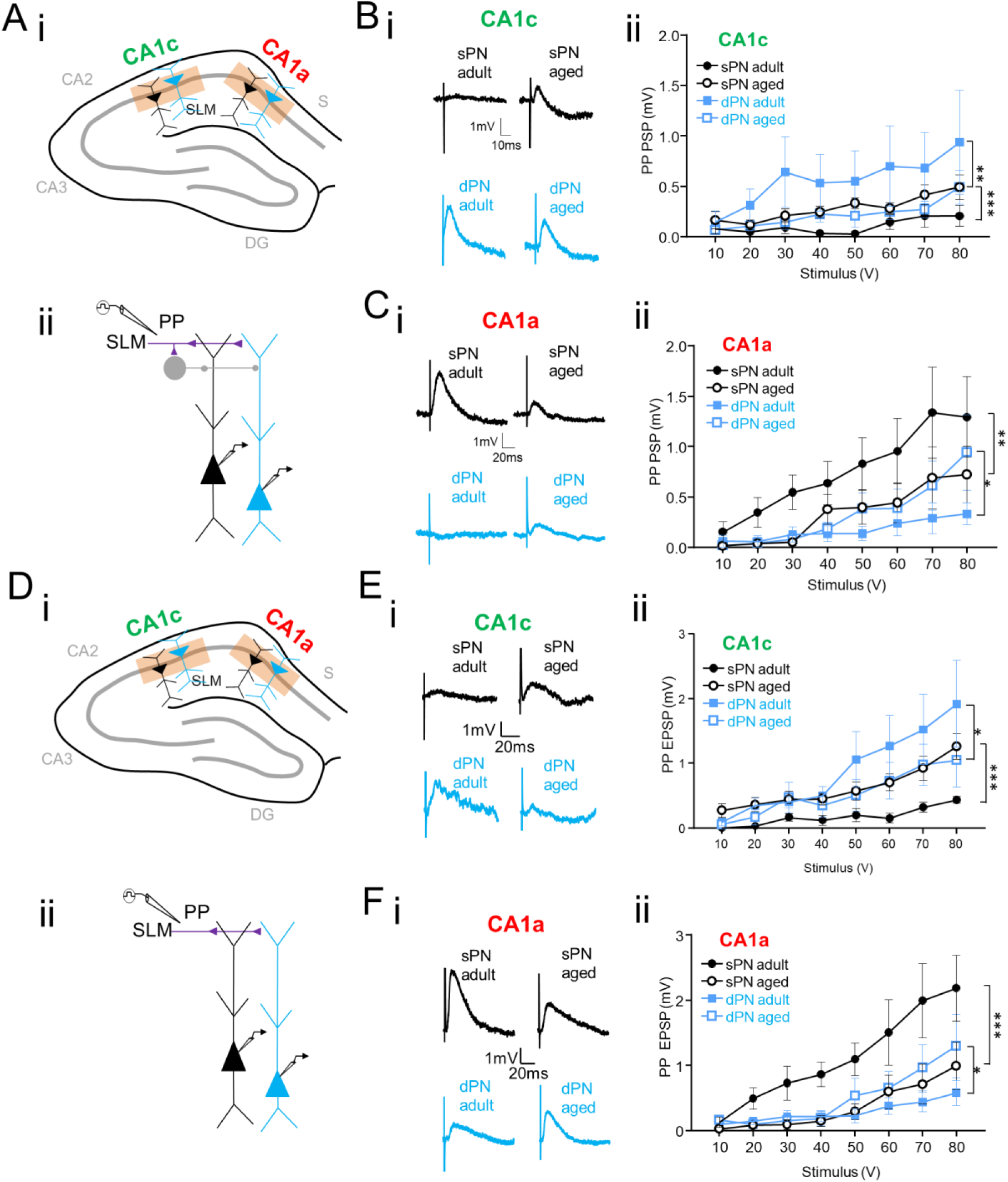
Aging equalizes perforant path excitation across transverse and radial axes. (A) In adult and aged male mice (i) whole cell recordings were targeted to superficial pyramidal neurons (sPNs, black) and deep PNs (dPNs, blue) in CA1c and CA1a of adult (3-6 month old) and aged (24-30 month old) mice. Perforant path (PP) responses were elicited by (ii) extracellular stimulation of stratum lacunosum moleculare (SLM) with inhibition intact. (B) (i) With inhibition intact, example PP postsynaptic potentials (PSPs) in CA1c (80V extracellular stimulation) and (ii) input-output curves of PP PSPs in sPNs/dPNs of CA1c across the two ages (aged: n = 8 from 5-6 mice/group; adult: n = 7 from 3-4 mice/group). (C) (i) With inhibition intact, example PP PSPs in CA1a (80V extracellular stimulation) and (ii) input-output curves of PP PSPs in sPNs/dPNs of CA1a across the two age groups (aged: n = 10-11 from 6 mice/group; adult: n = 10-11 from 6 mice/group). (D) Whole cell recordings were targeted to (i) sPNs and dPNs in CA1c and CA1a of adult and aged mice (ii) with an extracellular electrode place in SLM and with inhibition blocked. (E) (i) With inhibition blocked, example PP excitatory postsynaptic potentials (EPSPs) in CA1c (80V extracellular stimulation) and (ii) input-output curves of PP EPSPs in sPNs/dPNs of CA1c across the two ages (aged: n = 7-8 from 5-6 mice/group; adult: n = 7 from 3-4 mice/group). (F) (i) With inhibition blocked, example PP EPSPs in CA1a (80V extracellular stimulation) and (ii) input-output curves of PP EPSPs in sPNs/dPNs of CA1a across the two ages (aged: n = 10-11 from 6 mice/group; adult: n = 10-11 from 6-7 mice/group). For all above, error bars are +/-SE,*p < 0.05, **p < 0.01, ***p< 0.0001. Only statistical comparisons across age are indicated.

We next asked if aging altered the PP PSP by modulating the EPSP, IPSP or both. To examine the EPSP in isolation, we repeated our extracellular stimulation in SLM in the presence of GABA_A_ and GABA_B_ receptor antagonists (2 µM SR95531 and 1 µM CGP55845, respectively) (Figure 1Di,ii). In adult mice, the PP EPSPs displayed the same relationships across subpopulations as did the PP PSPs. Thus, the CA1c PP EPSP was larger in dPNs compared to sPNs (Figure 1E, p < 0.0001), and the CA1a PP EPSP was larger in sPNs compared to dPNs (Figure 1F, p < 0.0001), similar to our results in young mice (Masurkar et al., 2017). With aging, as with the PP PSP, these PP EPSP differences were eliminated through opposing effects on dPN and sPN EPSPs (Figure 1E, F). As such, in CA1c aging reduced the PP EPSP in dPNs while it increased the PP EPSP in sPNs, with the opposite effect in CA1a. These findings indicate that the observed PSP alterations are not associated with a change in inhibitory inputs, but represent purely a change in excitatory synaptic input. We found no alterations in the paired pulse ratio (PPR) measured in response to a pair of SLM stimuli, indicating a lack of change in presynaptic function (Supplemental Figure 1B). We also did not observe a change in the EPSP decay time constant (Supplemental Figure 1C), indicating a lack of change in passive or active dendritic integrative properties (Magee, 1998). Moreover, pharmacologic blockade of HCN channels (10 *μ*m ZD7288), which are enriched in the distal apical dendrite (Magee, 1998), did not impact PP EPSP magnitude relationships nor differentially impact decay time constants in CA1a sPNs across age groups (Supplemental Figure 2). Thus, the observed alterations must be due to changes in the strength of PP excitatory synaptic inputs onto PNs, presumably through changes in spine density, morphology, or glutamate receptor composition.

To determine if aging also modulated the synaptic connections CA1 received through its trisynaptic inputs, we next established input-output curves of SC-evoked PSPs via an extracellular electrode in SR (Figure 2Ai). We first examined the net SC PSP with inhibition intact (Figure 2Aii). In adult mice, the SC PSP was significantly larger in sPNs compared to dPNs in both CA1c and CA1a (Figure 2B, C; p = 0.0026 and p < 0.0001, respectively), similar to our findings in young mice (Masurkar et al., 2017). This difference likely results from the more marked feed-forward inhibition of dPNs exerted by parvalbumin-expressing interneurons, which preferentially target this population (Lee et al., 2014). Surprisingly, in contrast to the PP PSPs, aging had no effect on the SC PSPs in either sPNs or dPNs in either CA1c or CA1a (Figure 2B, C). We next recorded SC responses with inhibition blocked (Figure 2Di,ii) to determine if the isolated SC EPSPs followed the same pattern with aging as PP EPSPs. Prior studies that did not specify transverse or radial axis position have observed a reduction of SC excitation with aging (Barnes et al., 1992; Landfield et al., 1986; Deupree et al., 1993; Barnes et al., 1997; Barnes et al., 2000). Here, we found that in CA1c (Figure 2E) the SC EPSPs were similar in amplitude across the radial axis in adult mice, as in young mice (Masurkar et al., 2017), and that there was minimal change with aging. In CA1a (Figure 2F), SC EPSPs were also equivalent in amplitude in sPNs and dPNs, as in young mice (Masurkar et al., 2017). Yet in contrast to CA1c, with aging the CA1a SC EPSPs were reduced, and to a similar degree in both sPNs and dPNs. These results also suggest a compensatory reduction in feedforward inhibition in CA1a, both on sPNs and dPNs, since the SC PSPs did not show any change with aging in either population. With regard to the reduced SC EPSPs in CA1a, there were no clear differences in PPR (Supplemental Figure 1D) to suggest presynaptic dysfunction with aging as a contributing factor, consistent with prior studies (Barnes et al., 1997; Rosenzweig et al., 1997; Barnes et al., 2000). The EPSP decay time constant (Supplemental Figure 1E) also did not change to explain these alterations. These results also suggest that aging-related changes in SC EPSP responses relate to alterations in spine density, morphology, or receptor composition.

**Figure 2.**
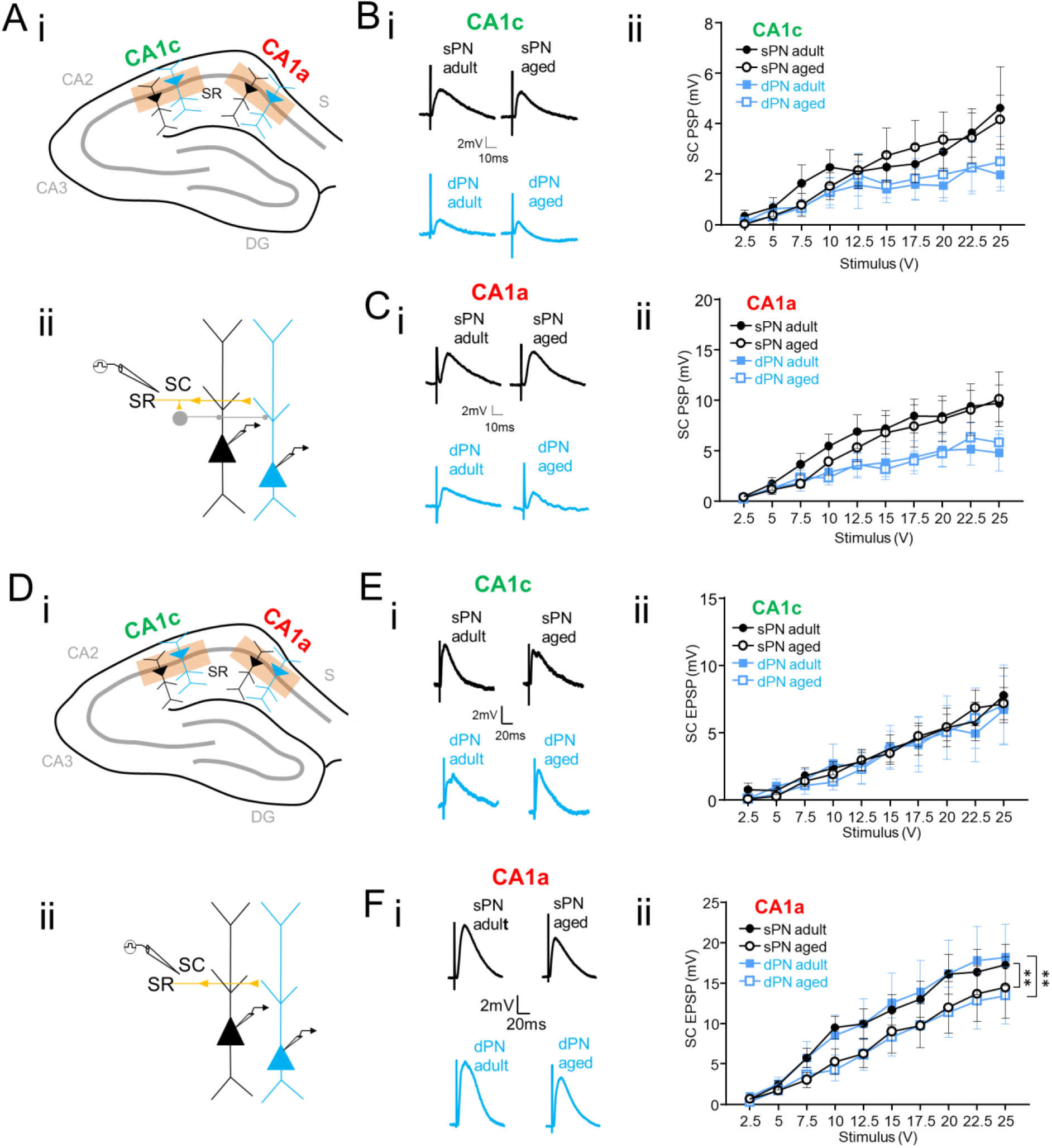
Aging has minimal impact on net Schaffer collateral excitation. (A) In adult and aged male mice (i) whole cell recordings were targeted to sPNs (black) and dPNs (blue) in CA1c and CA1a. Schaffer collateral (SC) responses were elicited by (ii) extracellular stimulation of stratum radiatum (SR) with inhibition intact. (B) (i) With inhibition intact, example SC PSPs in CA1c (25V extracellular stimulation) and (ii) input-output curves of SC PSPs in sPNs/dPNs of CA1c across the two ages (aged: n = 8 from 5-6 mice/group; adult: n = 7 from 3-4 mice/group). (C) (i) With inhibition intact, example SC PSPs in CA1a (25V extracellular stimulation) and (ii) input-output curves of SC PSPs in sPNs/dPNs of CA1a across the two ages (aged: n = 10 from 6-7 mice/group; adult: n = 9-10 from 5-6 mice/group). (D) In adult and aged male mice (i) whole cell recordings were targeted to sPNs and dPNs in CA1c and CA1a. Schaffer collateral (SC) responses were elicited by (ii) extracellular stimulation of stratum radiatum (SR) with inhibition blocked. (E) (i) With inhibition blocked, example SC EPSPs in CA1c (25V extracellular stimulation) and (ii) input-output curves of SC EPSPs in sPNs/dPNs of CA1c across the two age groups (aged: n = 8 from 5-6 mice/group; adult: n = 7 from 3-4 mice/group). (F) (i) With inhibition blocked, example SC EPSPs in CA1a (25V extracellular stimulation) and (ii) input-output curves of SC EPSPs in sPNs/dPNs of CA1a across the two ages (aged: n = 10 from 6-7 mice/group; adult: n = 9-10 from 6 mice/group). For all above, error bars are +/-SE, **p < 0.01. Only statistical comparisons across age are indicated.

In summary, aging appeared to affect PP- and SC-mediated excitation of CA1 PNs in distinct ways. Whereas the impact of aging on PP inputs varied according to transverse and radial position, irrespective of changes in inhibition, aging had no effect on the net SC-evoked PSP, although it did cause a selective reduction in the CA1a EPSP in both sPNs and dPNs. These data suggest the occurrence of a more profound, multipronged disruption in the organization of EC inputs to CA1 as a result of aging, compared to intrahippocampal CA3-CA1 connectivity.

### sPNs are differentially vulnerable to action potential-related excitability changes that amplify aging-related alterations in PP excitation

Our previous work on CA1 PNs in young (6-8 week) adult mice demonstrated that intrinsic excitability also varied based on somatic position along both transverse and radial axes, in a manner that enhances the functional specialization of these circuits established by their differential PP inputs (Masurkar et al., 2020). As such, we asked if any aging-related changes in intrinsic excitability of CA1 PNs were also dependent on location. Specifically, we assessed if such changes altered synaptic input processing to amplify or compensate for aging-related changes in synaptic responses uncovered above.

With regard to subthreshold properties measured at the soma (Supplementary Figure 3), we found that aging caused a ∼5.5 mV depolarization of the resting membrane potential V_rest_ in the sPNs of CA1a sPNs only, but otherwise had no significant effect on input resistance, membrane time constant, or sag ratio in either CA1c or CA1a that would influence the processing of excitatory synaptic responses. To examine suprathreshold properties, we provided increasing steps of input current (I) up to 450 pA, from a holding potential of -70 mV, and measured action potential firing (F) rates to generate F-I curves. In CA1c (Figure 3Bi, ii), aging produced a 15-20% increase in sPN firing rate. A small decrease in dPN firing rate did not reach statistical significance. In contrast, in CA1a (Figure 3Biii, iv), aging produced a reduction in both sPN and dPN firing, but with a particularly pronounced suppression in sPNs of nearly 3-fold at the F-I curve peak. In sum, in sPNs the aging-related alterations in firing rates are synergistic with the direction of PP EPSP change: increased in CA1c and decreased in CA1a. On the contrary, in dPNs there is either no statistically significant change (CA1c) or the change appears compensatory (reduced firing rate in CA1a in the face of increased PP EPSP).

**Figure 3.**
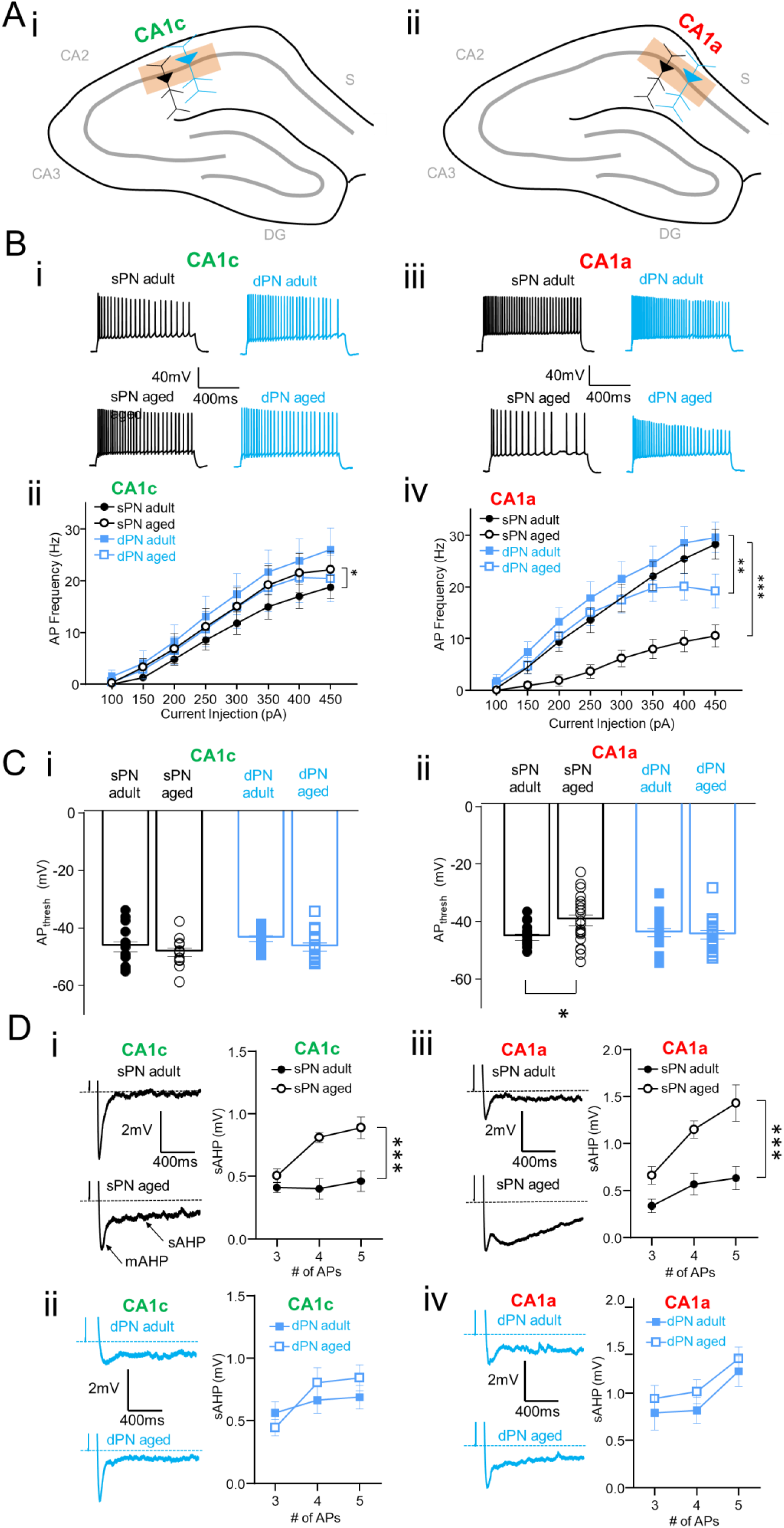
Aging preferentially impacts action potential-related excitability changes in sPNs. (A) In adult and aged male mice, whole cell recordings were targeted to sPNs and dPNs in (i) CA1c and (ii) CA1a. (B) (i) Example action potential (AP) firing elicited by 400 pA current injections in CA1c sPNs/dPNs of adult and aged male mice. (ii) Firing frequency-input current (F-I) curves for sPNs/dPNs in CA1c at both ages (aged: n = 12-14 from 8 mice/group; adult: n = 14-15 from 8-9 mice/group). (iii) Example AP firing (400 pA current injection) and (iv) F-I curves for CA1a sPNs/dPNs in both age groups (aged: n = 17-20 from 6-7 mice/group; adult: n = 17 from 10 mice/group). (C) In these same cells, AP threshold (AP_thresh_) for sPNs/dPNs in (i) CA1c and (ii) CA1a at both ages. (D) (i) Examples (left) of slow afterhyperpolarization (sAHP) (elicited by 5 APs, 5 trial average) in adult and aged CA1c sPNs. Indicated by arrows are the sAHP and medium AHP (mAHP). Plot (right) of sAHP amplitude as a function of APs in CA1a sPNs of both age groups (n = 8 from 5 mice/group). (ii) In adult and aged CA1c dPNs, examples (left) of sAHP and (right) sAHP amplitude as a function of APs (n = 8 from 3-4 mice/group). (iii) In adult and aged CA1a sPNs, examples (left) of sAHP and (right) sAHP amplitude as a function of APs (n = 7 from 3-4 mice/group). (iv) In adult and aged CA1a dPNs, examples (left) of sAHP and (right) sAHP amplitude as a function of APs (n = 7 from 3-4 mice/group). For all data above, error bars are +/-SE, *p < 0.05, **p < 0.001, ***p < 0.0001. Only statistical comparisons across age are indicated.

We then explored the mechanisms underlying these cell type-specific changes in firing rate. We first examined action potential (AP) threshold, as a prior study has demonstrated that aged CA1 PNs have altered voltage-gated sodium channel properties leading to a more positive AP threshold (Randall et al., 2012). In line with this previous finding, we observed a significant positive shift (> 6mV) in AP threshold in CA1a, however for sPNs only (Figure 3Ci,ii), consistent with the particularly pronounced reduction in AP firing in this population. Moreover, the magnitude of the AP threshold depolarization is greater than that of the V_rest_ in CA1a sPNs, thus likely to reduce AP firing rates even at their depolarized V_rest_. We next examined the effect of aging on the AP-evoked slow afterhyperpolarization (sAHP), which suppresses AP firing frequency. Moreover, the sAHP has been shown to increase with aging in CA1 PNs and associates with cognitive decline (Disterhoft et al., 1996; Landfield and Pitler, 1984; Tombaugh et al., 2005). For each PN subtype we induced the sAHP (Figure 3Di,iii; see Methods) after eliciting 3-5 APs, and then plotted the relation between sAHP magnitude and the number of APs. In both CA1c and CA1a sPNs, aging increased the size of the sAHP to a given number of APs, but had little impact on sAHP values in dPNs (Figure 3Di-iv). While the increased sAHP in CA1c sPNs lies in contrast with the slight increase in their firing rate with aging, in CA1a sPNs the concomitant effect of an increased sAHP with a depolarized AP threshold may contribute to their pronounced age-related firing rate reduction.

### CA1a sPNs in aged 3xTg-AD mice are selectively vulnerable to degeneration of PP inputs and reduction of SC drive

We next asked if the above age-related changes in PP synaptic responses and related processing may be exacerbated in the setting of AD pathology. We used the 3xTg-AD transgenic model which expresses both amyloidogenic and tauopathic genes (Oddo et al., 2003). In 3xTg-AD mice, amyloid plaques and tau-based neurofibrillary tangles are usually evident at ages above 12 months (Hirata-Fukae et al., 2008; Mastrangelo and Bowers, 2008; Oddo et al., 2003; Oh et al., 2010), although some variability in the timing of pathology onset has been suggested among different mouse colonies. Hence, we first verified the extent and localization of amyloid and tau pathology in mature animals from our own colony via 6E10 and AT8 antibody staining for amyloid plaques and neurofibrillary tangles, respectively (Supplementary Figure 4A-D). We subsequently focused our recordings in 3xTg-AD mice and age-matched wild type (WT) controls at 16-21 months, a time at which plaques and tangles were routinely observed. One important advantage in choosing this age is that this captures AD pathology in animals at a stage in the aging process when AD is prevalent in human patients. Furthermore, AD pathology in 3xTg-AD mice at this age has been reported to be predominant in CA1a (Oh et al., 2010), also as in human AD (Braak and Braak, 1997; Lace et al., 2009), a finding confirmed by our immunolabeling analysis (Supplementary Figure 4A-D). Thus, we limited our analysis to CA1a and focused on radial axis differences between wildtype and 3xTg-AD animals. Of note, AD transgene expression appears to be similar across the radial axis at this age, as evidenced by hAPP and HT7 antibody staining (Supplementary Figure 4E, F). Finally, experiments were performed using female mice only, as males in this line show a less robust phenotype (Clinton et al., 2007; Hirata-Fukae et al., 2008).

Given that we observed above an impact of aging specifically on the PP EPSP (Figure 1D-F), we first asked if the presence of concomitant AD pathology exacerbated this effect. As such, using the same slice preparation method applied to aged mice, we recorded PP EPSPs via extracellular stimulation in SLM in the presence of GABA blockers (Figure 4A). Importantly, the magnitude of the PP EPSPs in aged WT mice were similar in sPNs and dPNs, recapitulating our findings in aged C57Bl/6J mice (Figure 2C). Moreover, compared to aged WT mice, there was a significant decrease in the PP EPSP in sPNs of aged 3xTg-AD mice (Figure 4B), presumably exacerbating the normal effect of aging. In contrast, there was no difference between aged WT and transgenic mice in the PP EPSPs recorded in dPNs (p = 0.89). This resulted in significantly smaller PP EPSPs in sPNs versus dPNs in the aged 3xTg-AD mice (p = 0.0001), with an over 70% difference at the peak of the stimulus-response curve. This selective change in sPNs was not explained by alterations in markers of presynaptic efficacy or postsynaptic dendritic excitability (Supplemental Figure 5B,C).

**Figure 4.**
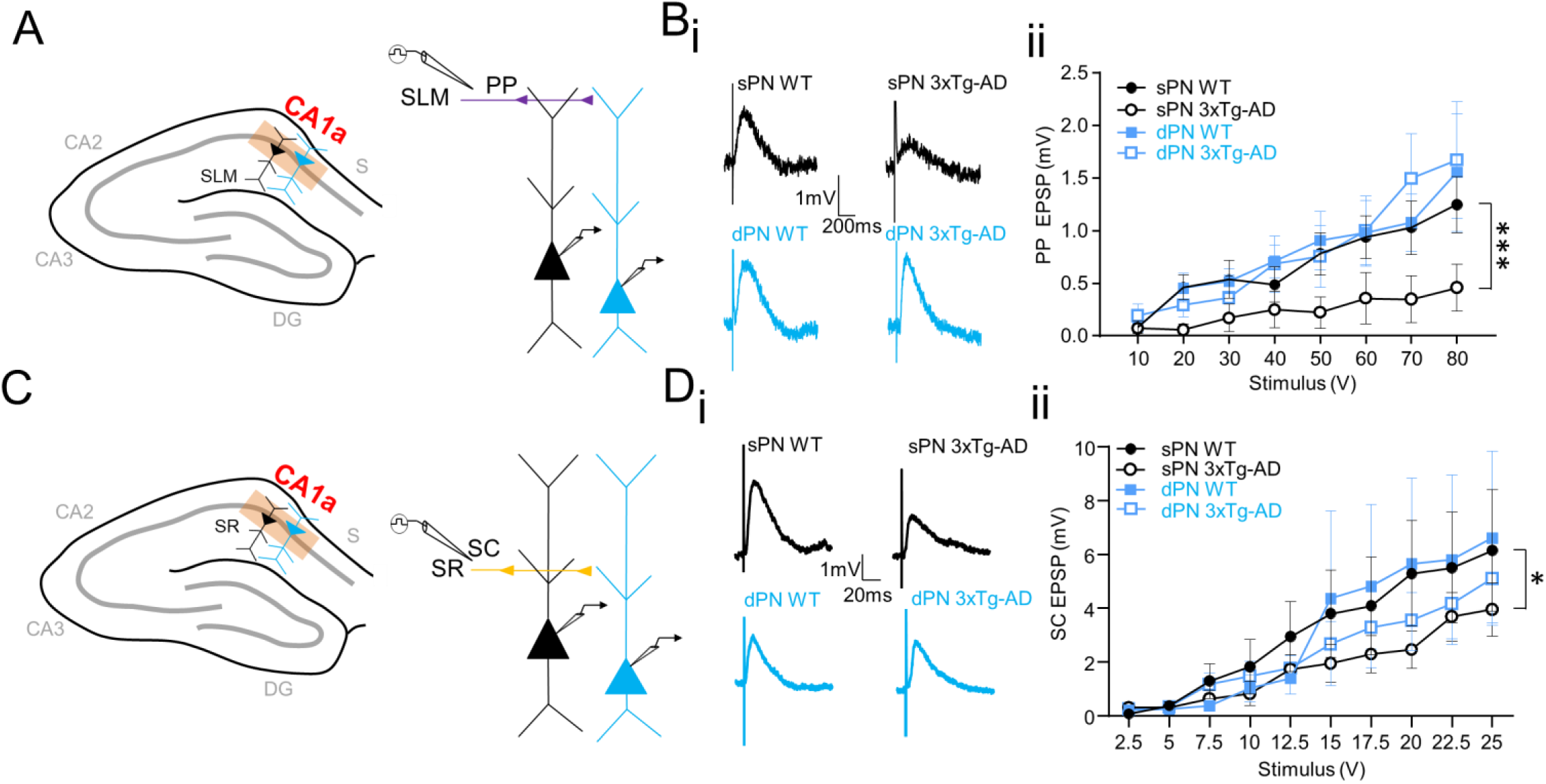
Selective reduction of PP and SC EPSPs in CA1a sPNs of aged 3xTg-AD mice. (A) Whole cell recordings targeted to (left) CA1a sPNs/dPNs in CA1a of female 3xTg-AD and WT mice (16-21 months old) with (right) PP EPSPs elicited by extracellular stimulation of SLM with inhibition blocked. (B) (i) Example PP EPSPs in CA1a (80V stimulation). (ii) input-output curves of PP EPSPs in CA1a sPNs/dPNs of 3xTg-AD versus WT mice (AD: n = 8-11 from 5 mice/group; WT: n = 10-11 from 5 mice/group). (C) Whole cell recordings were targeted to (left) CA1a sPNs/dPNs with (right) SC EPSPs elicited by extracellular stimulation of SR with inhibition blocked. (D) (i) Example SC EPSPs in CA1a (25V stimulation). (ii) input-output curves of SC EPSPs in CA1a sPNs/dPNs of 3xTg-AD versus WT mice (AD: n = 8-11 from 5 mice/group; WT: n = 10-11 from 5 mice/group), *p < 0.05; ***p < 0.0001. Only statistical comparisons across genotype are indicated.

We next asked if SC EPSPs were also differentially reduced in CA1a sPNs, or underwent equivalent reduction in CA1a sPNs and dPNs as we observed with aging alone. We used extracellular stimulation of SR in the presence of GABA blockers (Figure 4C), as described earlier, to record SC EPSPs. As with PP EPSPs, we observed that AD transgene expression led to a selective decrease in the magnitude of the SC EPSP in sPNs compared to that seen in aged WT mice (Figure 4D). Although there was a trend for AD transgene expression to decrease the SC EPSP in dPNs, this was not statistically significant. As with the PP EPSPs, there was no selective change in PPR or EPSP decay time constant to explain the reduction of SC EPSP magnitude in sPNs (Supplemental Figure 5D, E).

To determine whether the reduction of PP and SC EPSPs in sPNs could be explained by a selective loss of dendritic spines, we dialyzed sPNs and dPNs during patch clamp recordings with an intracellular solution containing 25*μ*m AlexaFluor 594. We then used two-photon laser scanning microscopy to visualize and quantify spine density in the distal and proximal apical dendrites that are targeted by PP and SC inputs, respectively (Figure 5A; see Methods). In aged WT mice, we first noted that the distal dendrite spine density was nearly two-fold larger in CA1a sPNs compared to dPNs (Figure 5Bi, ii) despite their equivalent PP EPSP magnitudes (Figure 4Bi,ii). This spine density ratio is similar to what we have observed in CA1a of young adult C57BL/6J mice, at an age when sPNs feature larger PP EPSPs than dPNs that is largely explained by such spine density differences (Masurkar et al., 2017). This suggests that aging-related changes in CA1a responses delineated earlier (Figure 1Fi, ii) are not related to changes in spine density. We additionally observed that the distal spine density in sPNs was reduced greater than 50% in aged 3xTg-AD compared to aged WT mice (Figure 5Bi, ii), concordant with the decline in the PP EPSP magnitude (Figure 4Bi, ii). In contrast, there was no difference in distal dendrite spine density between genotypes in dPNs. Thus, the selective loss of distal dendritic spines in sPNs provides a likely mechanism to explain, at least in part, the reduced PP EPSP magnitude in aged 3xTg-AD mice. In contrast, there was no evidence of selective spine loss in the proximal dendrite of sPNs to explain the reduced SC EPSP magnitude in age 3xTg-AD mice (Figure 5Ci,ii). This implies that the reduced SC excitatory response in sPNs of aged 3xTg-AD mice is solely due to postsynaptic mechanisms other than spine loss.

**Figure 5.**
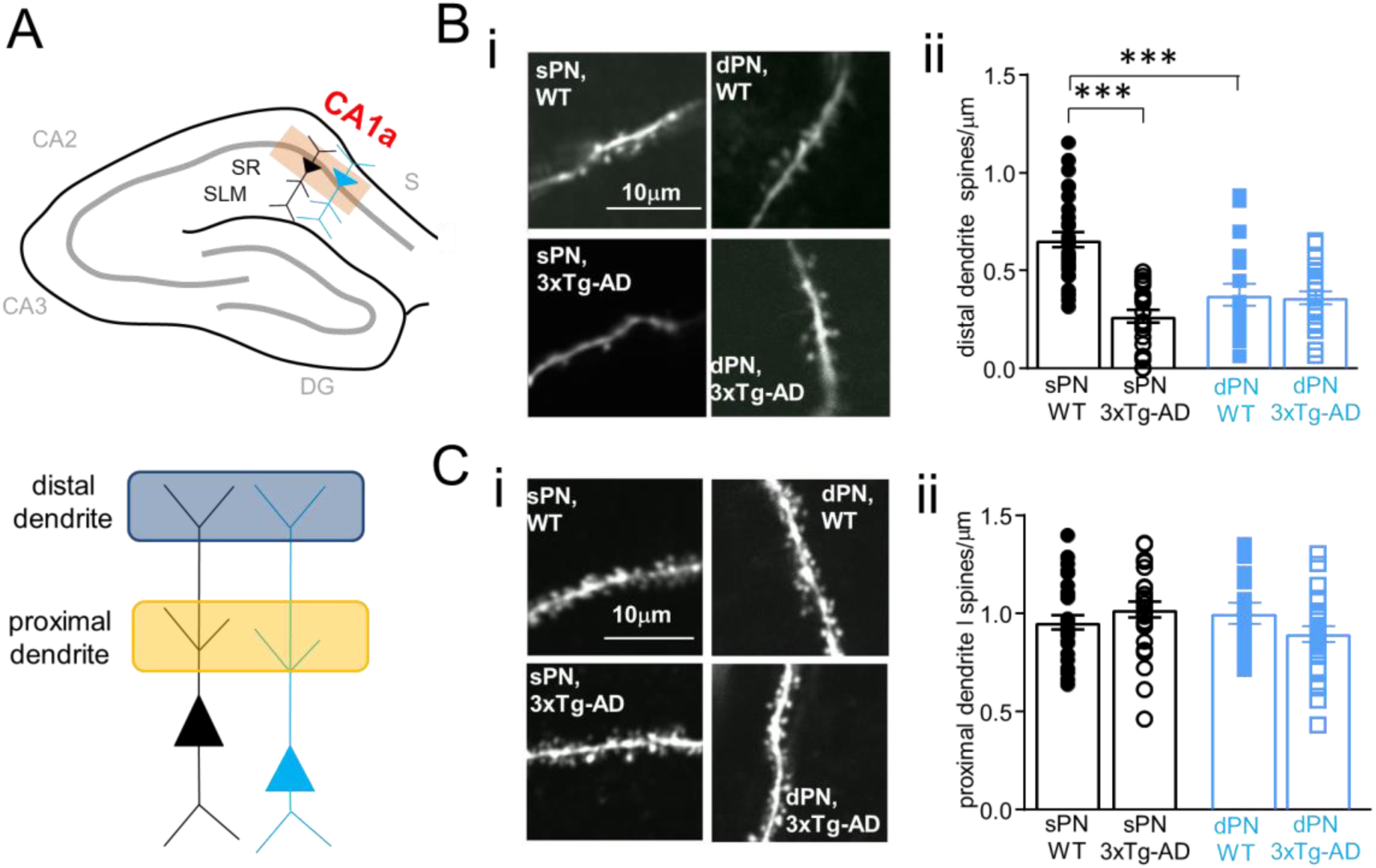
Selective reduction of distal dendrite spine density in CA1a sPNs of aged 3xTg-AD mice. (A) In aged female 3xTg-AD and WT mice, CA1a sPNs and dPNs (above) were dialyzed with 25*μ*m AlexaFluor 594 via the patch pipette in order to visualize (below) spines in the distal and proximal apical dendrite via 2-photon microscopy. (B) Across radial axis and genotype, (i) example images of spines visualized via 2-photon microscopy on distal apical dendrite branch segments and (ii) spine density (AD: n = 24-26 segments over 4-5 cells from 4-5 animals/group; WT: n = 20-33 segments over 4-5 cells from 4-5 animals/group). (C) Across radial axis and genotype, (i) example images of spines visualized via 2-photon microscopy on proximal apical dendrite branch segments and (ii) spine density (AD: n = 25-27 segments over 4-5 cells from 4-5 animals/group; WT: n = 16-32 segments over 4-5 cells from 4-5 animals/group). For all above, error bars are +/-SE. For all above, error bars are +/-SE, ***p < 0.0001

### CA1a sPNs in aged 3xTg-AD mice are differentially vulnerable to AP-related excitability changes that amplify reduced excitatory drive

Given our finding that aging results in specific changes in PN excitability, we finally explored if intrinsic excitability changes in aged 3xTg-AD mice may also potentially influence synaptic input processing to exacerbate the reduced synaptic drive of CA1a sPNs. With regard to subthreshold properties, we found no difference in resting potential, input resistance, membrane time constant or sag ratio between aged WT and 3xTg-AD mice (Supplementary Figure 6B-E). Examining suprathreshold properties, we observed a surprising increase in AP firing in dPNs of aged 3xTg-AD compared to aged WT mice, reaching over 40% at peak of the F-I curve (Figure 6B). In contrast, there was a slight decrease in AP firing in sPNs of 3xTg-AD mice, though not statistically significant. As a result, in aged 3xTg-AD mice the dPNs had a significantly higher rate of AP firing compared to sPNs (p = 0.0005), about 2-fold at the peak of the F-I curve, not unlike what was observed in aging. We further observed a statistically significant increase in the sAHP amplitude of sPNs in aged 3xTg-AD compared to WT mice (Figure 6Ci). This could contribute to the observed trend for a reduced firing rate of sPNs, although this did not reach statistical significance in the sample examined. In contrast, in dPNs there were no changes in sAHP (Figure 6Cii) or AP threshold (Supplementary Figure 6F) across genotype that would drive their increased firing rates. Thus, like aging, AD impacts AP firing in CA1a to result in significantly higher firing rates in dPNs relative to sPNs, albeit via distinct mechanisms. Moreover, similar to aging, this differential intrinsic excitability change would amplify the effect on action potential output of a biased reduction in excitatory synaptic response magnitude in sPNs compared to dPNs.

**Figure 6.**
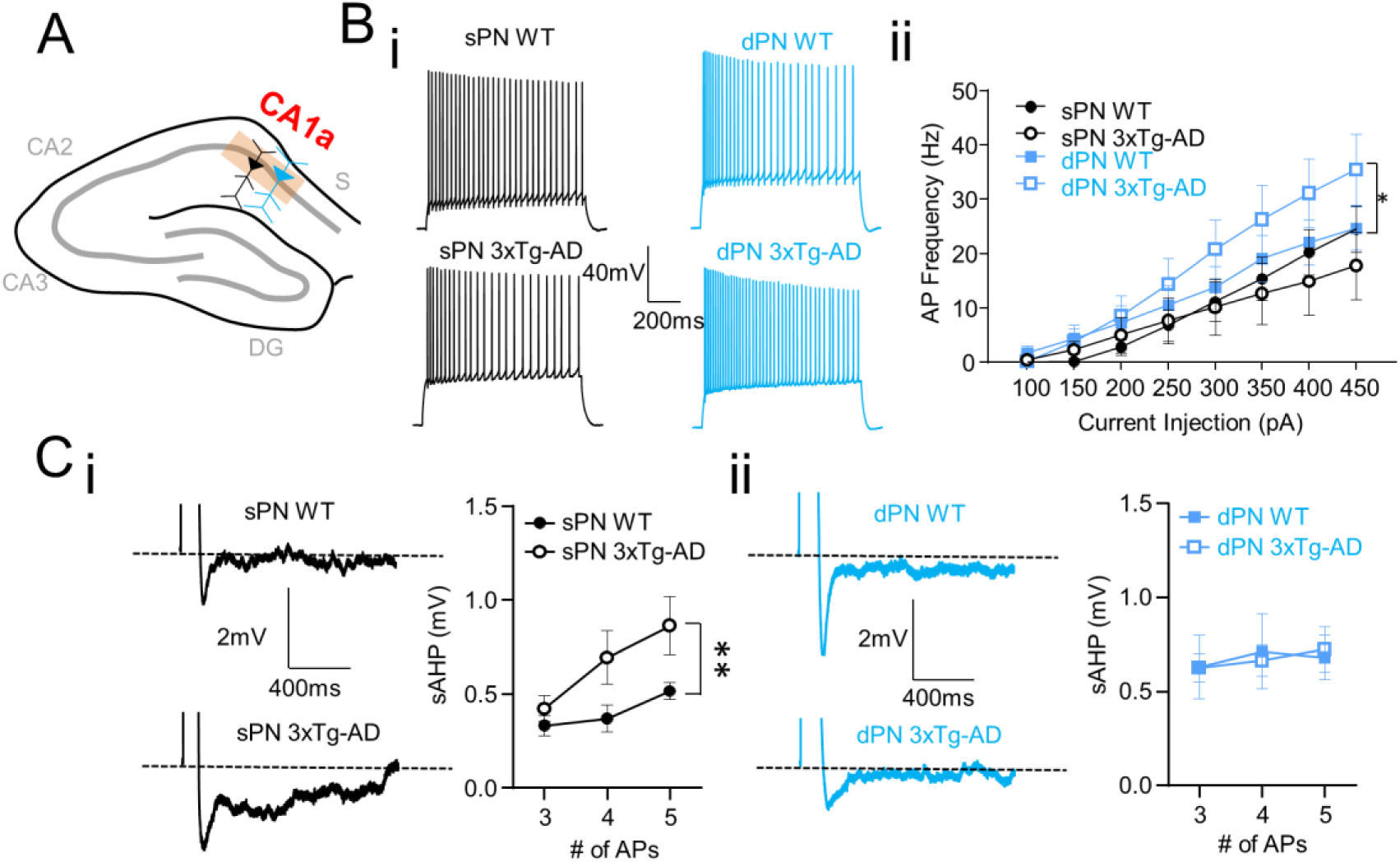
Differential susceptibility of CA1a sPNs to AP-related excitability changes in aged 3xTg-AD mice. (A) Whole cell recordings of CA1a sPNs and dPNs were performed in aged female 3xTg-AD and WT mice to evaluate AP-related excitability properties. (B) Across radial axis and genotype, (i) examples of AP firing (450 pA current injection) and (ii) F-I curves (AD: n = 8-11 from 5 mice/group; WT: n = 10-11 from 5 mice/group). (C) (i) In sPNs across genotype, examples (left) of sAHP (induced by 5 APs, average of >= 5 trials) and plots (right) of sAHP as a function of APs (n = 7-9 from 5 mice/group). (ii) In dPNs across genotype, examples (left) of sAHP (induced by 5 APs, average of >= 5 trials) and plots (right) of sAHP as a function of APs (n = 11 from 5 mice/group). Error bars are +/-SE, *p < 0.05; **p < 0.01.

## DISCUSSION

In this study, we observed that the effect of aging and AD on excitatory drive and intrinsic excitability of CA1 PNs depends on somatic location. Notably, we revealed that, compared to SC input and intrinsic excitability, aging and AD pathology have unique and strong effects on PP excitation, summarized in Figure 7. In non-aged adult mice, PP impact normally varies depending on CA1 PN position across the radial and transverse axes, shaping the functional subspecialization of CA1 PNs (Masurkar et al., 2017). In aged mice, such differences are eliminated through opposing reductions and increases in PP impact depending on CA1 PN position (Figure 7AB). This results in a uniform PP impact across CA1 at an intermediate magnitude. Moreover, with concurrent AD pathology in aged 3xTg-AD mice primarily concentrated in CA1a (Figure 7C), such changes are exacerbated, with a particularly profound reduction in PP inputs in CA1a sPNs (Figure 7D). These results illustrate a novel feature of how aging and AD can disrupt excitatory flow through the hippocampal circuit.

**Figure 7.**
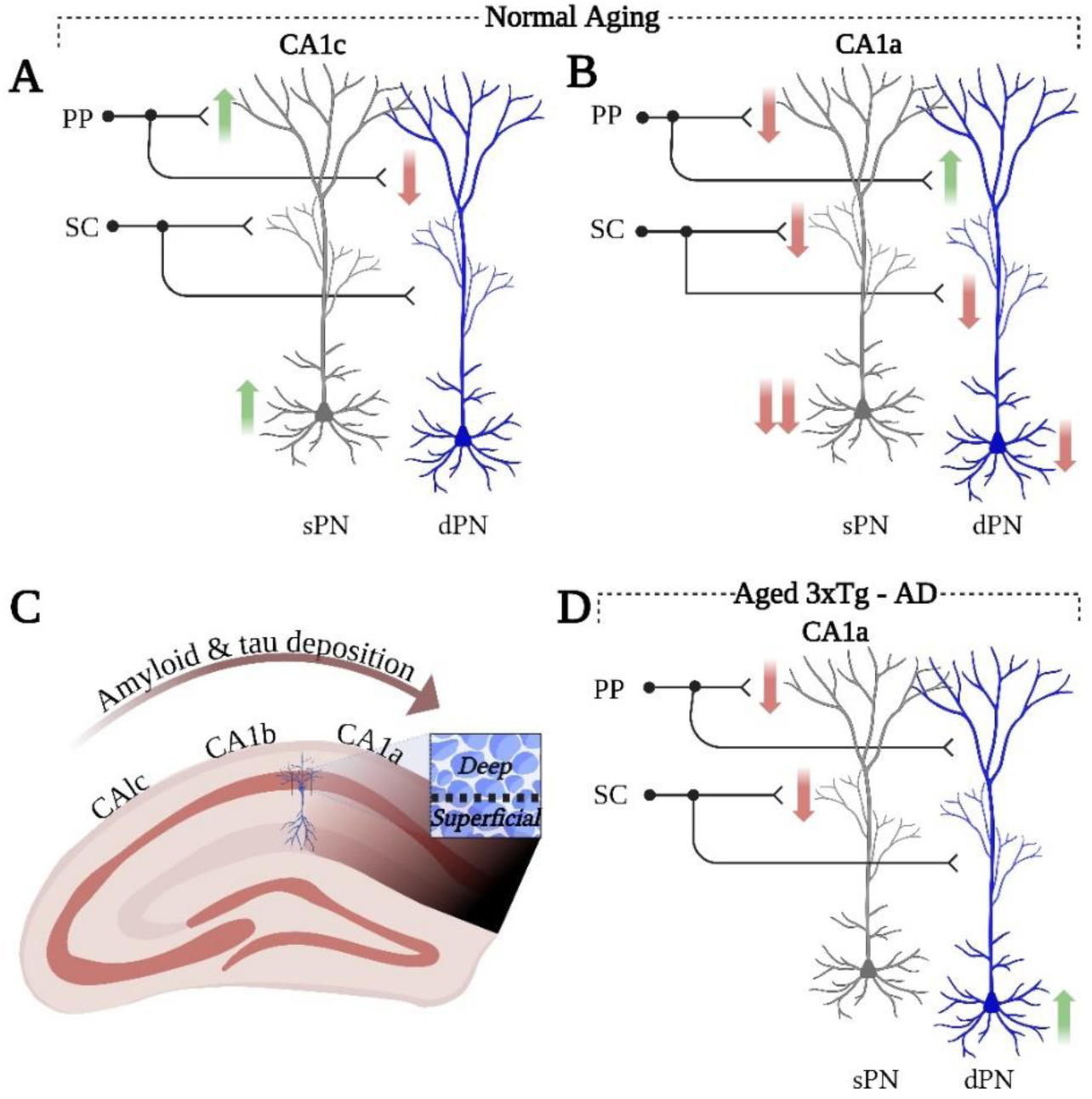
Summary of aging- and AD-related changes to EC→CA1 excitatory flow. Schematic showing aging-related changes in PP and SC excitatory impact and intrinsic excitability measured at the soma in sPNs and dPNs of (A) CA1c and (B) CA1a. In aged 3xTg-AD mice with (C) highest burden of AD pathology in CA1a, (D) scheme showing AD-related changes in PP and SC excitatory impact and intrinsic excitability in sPNs and dPNs of CA1a. Up arrow indicated increased synaptic impact or excitability; down arrow indicates decreased synaptic impact of excitability. Images created in Biorender.

### Mechanisms underlying aging-related equalization of PP input responses across the radial axis

Our results suggest that aging impacts PP excitatory responses with opposing effects on dPNs and sPNs that depend on transverse axis position. In contrast, aging-related changes to SC excitatory responses depended only on transverse axis position (reduced responses in CA1a). This further highlights the distinction between these two pathways that both synapse on the CA1 PN apical dendrite, albeit on distinct compartments: PP inputs on the distal dendrite and SC inputs on the proximal dendrite.

With regard to specific mechanisms, we note no change in PP presynaptic function as measured by the PPR. This contrasts with aging-related changes to the EC→DG pathway, which undergoes a strong reduction in presynaptic integrity and function (Froc et al., 2003; Yassa et al., 2011; Amani et al., 2021). One explanation may be that the cortical efferents to these hippocampal subregions derive from functionally and architecturally separate lamina (van der Linden and Lopes da Silva 1998; Gloveli et al., 1999; Tahvildari and Alonso 2005), with EC layer II as the primary source to DG and EC layer III as the dominant input to CA1 (van Groen et al., 2002). With respect to postsynaptic mechanisms, there has been limited examination of the impact of aging on dendritic ion channels that shape dendritic responses to excitatory input, with one study finding an aging-related reduction of A-type potassium current as measured at the soma (Alshuaib et al., 2001). Here we find that decay time constant of the EPSP, which is influenced by dendritic excitability, was unchanged with aging in dPNs and sPNs, thereby not consistent with the alteration in cortical response amplitudes. Moreover, our spine imaging of aged WT mice revealed a distal dendrite spine density in CA1a nearly two-fold larger in sPNs compared to dPNs. This difference in spine density is similar to what we have observed in CA1a of young adult mice, an age at which PP EPSPs are larger in sPNs than dPNs in proportion to the increased spine density (Masurkar et al., 2017). In sum, these findings point to mechanisms at the level of spine morphology subtype distribution, glutamate receptors, or other molecular aspects of postsynaptic structure. Prior studies of the CA1 SLM region in aged rodents have not revealed any changes in the density of mushroom and thin spines, the number of perforated and nonperforated synapses, or the density of AMPA and NMDA receptors within these synapses (Xu et al., 2018; Buss et al., 2021). However, this has not been addressed specifically across radial and transverse PN subpopulations.

### Identity of PP inputs driving aging-related response changes and relevance to behavior

In young adult mice, we previously found that PP responses in CA1a were significantly larger in sPNs versus dPNs, whereas in CA1c this was reversed, with larger PP responses in dPNs versus sPNs (Masurkar et al., 2017). We additionally found that in CA1a this difference was largely mediated by preferential LEC drive of sPNs, with a minor contribution of MEC input. In contrast, in CA1c this was mediated by preferential MEC drive of dPNs, and a minor contribution of LEC input. In this study, we find that with aging, such differences in PP responses across radial and transverse axes are eliminated by the following changes: in CA1a, PP responses are reduced in sPNs and are increased in dPNs; in CA1c, PP responses in are reduced in dPNs and increased in sPNs. What could be the efferent source of the aging-related increase in PP EPSPs in CA1a dPNs and CA1c sPNs? In either case, it may be driven by a heightened response to LEC, MEC, or both inputs. In contrast, we anticipate that the aging-related ∼50% reductions of the PP EPSP in CA1a sPNs and CA1c dPNs are largely due to reduced LEC and MEC responses, respectively, because our previous study in young adult mice suggested that these inputs drive the majority (>80%) of the total PP EPSP in these PNs (Masurkar et al., 2017). Consistent with our proposed mechanism above, we postulate that spines specifically targeted by LEC or MEC undergo aging-related remodeling to elicit these changes. Of note, LEC→CA1 and MEC→CA1 synapses show ultrastructural differences (Bloss et al., 2018) that may relate to aging-related changes on specific PNs.

Elucidating the efferent identity corresponding to the PP responses may shed led light on aging-related memory deficits. Spatial memory dysfunction with aging associates with a reduction in place field specificity (Barnes et al., 1983; Wilson et al., 2005; Yan et al., 2003), which is established by MEC layer III input to CA1 PNs (Brun et al., 2008). While aging-related changes of MEC itself certainly could drive this dysfunction, altered integration of this input across CA1 dPNs and sPNs, both critical for spatial memory and different aspects of spatial coding (Mizuseki et al., 2011; Danielson et al., 2016; Maroso et al., 2016, Geiller et al., 2017, Sharif et al., 2021), may also contribute significantly to this deficit. Analogously, altered response to LEC input by CA1 PNs may relate to aging-related non-spatial memory deficits (Hendricks et al., 2021; LaSarge et al., 2007; Patel and Larson, 2009; Prediger et al., 2006; Roman et al., 1996; Soontornniyomkij et al., 2012; Terranova et al., 1994). Moreover, such postulated changes in MEC and LEC responses are additionally amplified by PN-specific alterations in SC impact and intrinsic excitability, most notably in CA1a, further contributing to a reshaping of CA1 population coding that may negatively impact memory-guided behavior.

### Mechanisms of selective AD-related distal dendrite spine loss in CA1a sPNs

In 3xTg-AD mice we find that reduced PP responses in CA1a sPNs correlate with a reduced spine density in sPN distal apical dendrite, the site of PP synapses. In contrast, the proximal apical dendrite, the site of SC synapses, did not feature spine loss. In this transgenic model, while they did not specify which hippocampal subregion or PN dendritic compartment, Bittner and colleagues observed that hippocampal spine loss is evident between 15-20 months of age, similar to mice in our study, and is associated with proximity to amyloid plaques or intra-dendritic accumulation of amyloid and phosphorylated tau (Bittner et al., 2010). This would suggest that, compared to the proximal compartment, the distal dendrites of sPNs either are differentially susceptible to degenerative effects of extracellular amyloid plaques, are more prone to develop intra-dendritic amyloid and phosphorylated tau, or are more vulnerable to the toxic effects of this intradendritic dual pathology. Moreover, sPNs are more susceptible to this effect than dPNs. Of note, sPNs display increased expression of the 5-HT1A receptor (Cembrowski et al., 2016), which promotes amyloid-induced tau phosphorylation (Wang et al., 2016), and c-Jun N-terminal kinases (Maroso et al, 2016), which facilitate tau phosphorylation and amyloid-induced spine loss (Coulson et al., 2006; Yarza et al., 2016). Lastly, we find that sPNs in this model show increases in the calcium-dependent sAHP, which suggests that they may be more susceptible to calcium dyshomeostasis, which can drive spine loss (Wang and Zheng, 2019).

### Behavioral impact of AD-related changes on PP inputs in CA1a

Memory deficits in AD models may reflect the impact of pathology across the EC→CA1 network. Spatial memory deficits in AD models feature abnormalities in various features of place cells (Ciupek et al., 2015; Mably et al., 2017; Zhao et al., 2014), although a major driver of this change may be more attributed to MEC dysfunction (Fu et al., 2017; Jun et al., 2020; Ness and Schultz, 2021), given the relative absence of pathology in CA1c. In contrast, with regard to non-spatial memory, 3xTg-AD mice develop deficits in novel object recognition and olfactory behavior well after EC involvement, at an age that correlates better with the emergence of prominent CA1a pathology (Arsenault et al., 2011; Cantarella et al., 2015; Cassano et al., 2011; Clinton et al., 2007; Coronas-Samano et al., 2014; Mastrangelo and Bowers, 2008; Mitrano et al., 2021; Stover et al., 2015). This suggests that these deficits result from the concomitant impact of pathology on LEC and CA1a circuits that specifically integrate LEC input, notably sPN-driven circuits (Masurkar et al., 2017; Li et al., 2017). Here we find that in aged 3xTg-AD mice, PP drive is severely compromised selectively in sPNs. This reduced drive, most likely related to the LEC response (Masurkar et al., 2017), and which are amplified from those found in aging alone, may serve as a critical contributor to these behavioral deficits. Processing of LEC input by sPNs is likely further compromised by the significantly reduced SC impact on these neurons. This hypothesis would further assume that CA1a dPNs cannot effectively compensate, despite uncompromised PP impact, minimally reduced SC responses, and increased action potential firing capability. Of note, human AD features deficits in olfactory function, including odor memory, that precede spatial memory loss (Masurkar and Devanand, 2014). This arises at an early stage when AD pathology extends from LEC to CA1a, at which there is morphological compromise in a subset of distal dendrites within SLM (Braak and Braak, 1997; Lace et al., 2009; Masurkar and Devanand, 2014). It remains to be explored whether these vulnerable distal dendrites in human CA1 originate from sPNs.

## METHODS

Experiments were performed according to National Institutes of Health guidelines and with approval from the Columbia University and New York University School of Medicine Institutional Animal Care and Use Committee.

### Animals

Aging experiments were performed on adult (3-6 month) and aged (24-30 month) C57BL/6J mice (Jackson Labs stock # 000664). We restricted our analysis to males to enable comparison to previous work on these neuronal subpopulations (Masurkar et al., 2017; Masurkar et al., 2020). For AD experiments, we used 3xTg-AD mice (Jackson Labs stock # 004807) and WT controls (C57BL/6J:129SvEvTac hybrid) at 16-21 months old. We restricted our analysis to females due to the known minimized phenotype in males of this line (Clinton et al., 2007; Hirata-Fukae et al., 2008). AD experiments were performed by individuals blinded to genotype.

### Slice Preparation

Mice were anesthetized with isoflurane and subsequently underwent intracardiac perfusion with cooled dissection artificial cerebrospinal fluid (dACSF). The dACSF was comprised of 10 mM NaCl, 25 mM NaHCO_3_, 2.5 mM KCl, 1.5 mM NaH_2_PO_4_, 7 mM MgCl_2_, 0.5 mM CaCl_2_, 195 mM sucrose, 10 mM D-glucose, and 2 mM Na-pyruvate, which was brought to pH 7.3 with carbogen (95%/5% O_2_/CO_2_). After decapitation, hippocampi were dissected in cooled, carbogenated dACSF and affixed to a 4% agar block. The block was transferred to a vibratome (Leica VT1200S) filled with cooled, carbogenated dACSF. Temperature was maintained below 5°C. Beginning at the dorsal end, but discarding the first two slices, six to seven consecutive 400-*μ*m transverse hippocampal slices were sectioned bilaterally. These slices were transferred to an incubation chamber at 34°C containing a carbogenated 50%/50% mixture of dACSF and recording ACSF (rACSF). The rACSF was comprised of 125 mM NaCl, 25 mM NaHCO_3_, 2.5 mM KCl, 1.25 mM NaH_2_PO_4_, 1 mM MgCl_2_, 2 mM CaCl_2_, 22.5 mM D-glucose, 3 mM Na-pyruvate, and 1mM L-ascorbic acid, brought to pH 7.3 with 95%/5% O /CO . After 25 minutes at 34°C, the temperature was reduced to 20-25°C. Slices remained in the incubation chamber until used for experiments.

### In Vitro Electrophysiology

Slices were placed in a chamber with carbogenated rACSF maintained at 34 °C. CA1 PNs were visualized using a 60x, 0.9 NA water immersion objective, differential interference contrast imaging, and an infrared camera (Olympus) that interfaced with ImageJ. Before adding blockers of inhibition, a cut was placed at CA3/CA2 to prevent epilepsy. Patch pipettes were fashioned from fire-polished borosilicate glass pulled to 3.8-5.0 M*Ω* (Sutter) and were back filled with intracellular solution comprised of 135 mM KMeSO_3_, 5 mM KCl, 2 mM NaCl, 0.2 mM EGTA, 10 mM HEPES, 10mM Na_2_phosphocreatine, 5 mM MgATP, 0.4 mM Na_2_GTP, 25 *μ*M Alexa Fluor 594 (Thermo Fisher Scientific), with pH adjusted to 7.3. Electrophysiological measurements were achieved in the current clamp mode of the Multiclamp 700B amplifier using pClamp 9 software (Axon Instruments) and a Digidata 1322A digitizer. When appropriate, the GABA receptor blockers SR95531 and CGP55845 (Sigma), or HCN channel blocker ZD7288 (Tocris), were added to the bath, allowing 5-10 minutes to take full effect. For targeted whole-cell recording, sPNs were identified as the first 1-2 layers of neurons in stratum pyramidale below the stratum radiatum border. The dPNs comprised the subsequent layers extending to the stratum oriens border. PNs were targeted in alternating fashion, as much as possible, across the radial axis and also the transverse axis when relevant. Upon break-in, resting membrane potential was immediately noted and recorded. After 2-5 minutes of dialysis with intracellular solution, protocols were run to assay intrinsic and synaptic excitability, while maintaining membrane potential at -70 mV. PN identity was confirmed by an accommodating action potential firing pattern, presence of voltage sag to hyperpolarization, and resting membrane potential below -60mV. Experiments were terminated if resting membrane potential depolarized by more than 5mV or if series resistance increased above 25 M*Ω.*

Intrinsic excitability measurements were evaluated as follows. V_rest_ was noted immediately after gigaseal rupture, and the remaining properties were measured after 2-5 minutes. Other than the sAHP, these properties were measured while maintaining V_m_ at -70mV. R_in_ was derived from the average steady state voltage response to a -10 pA, 500 ms current injection over 10 trials. The *τ*_m_ was computed from the same voltage response, by fitting a monoexponential to the initial membrane hyperpolarization. Sag was induced by injecting negative current, of 1 s duration, enough to hyperpolarize membrane potential V_initial_ to a V_sag_ of -100mV to -105mV, which then depolarized to V_end_ before the end of the square pulse. Sag ratio was calculated as (V_end_-V_sag_)/(V_initial_-V_sag_). F-I curves were measured by injecting depolarizing current injections of 1 s duration and increasing amplitude, and quantifying the number of APs elicited. AP threshold was derived from the first AP generated in the F-I-curve. For sAHP-AP curves, the sAHP was measured at a V_m_ of -60mV. To induce the sAHP, a 100 ms depolarizing current injection was provided, enough to elicit 3, 4, or 5 APs. For each AP number, at least 5 trials were averaged. This protocol induced a medium AHP (mAHP), peaking less than 100ms after the current injection, followed by the sAHP which was measured at its peak at around ∼150-450ms after the current injection.

For synaptic responses, low resistance stimulating electrodes were fashioned out of fire-polished borosilicate glass (Sutter), backfilled with rACSF, and placed in stratum lacunosum moleculare and stratum radiatum. Voltage pulses (0.1ms duration, every 30s) were delivered to the stimulating electrode via a constant voltage isolator (Digitimer Ltd). For the paired pulse ratio (PPR), two extracellular stimuli were given at 20Hz to give two EPSPs The PPR was calculated by dividing the amplitude of the second EPSP by that of the first EPSP. The tau_decay_ of EPSPs were computed by fitting a monoexponential to the downward phase of the potential.

### Two-Photon Imaging

CA1 PNs were dialyzed with 25 *μ*M Alexa Fluor 594 via the patch pipette. Spine imaging was initiated at least 35 minutes after break-in to allow filling of the distal apical dendrites. Only one PN per slice was analyzed to avoid intermingling of dendrites from different PNs. Imaging was performed by an individual who was blinded to both the genotype and the superficial/deep identity of the PN, using a Prairie Technologies Ultima two-photon microscope and an excitation wavelength of 820 nm. Proximal apical dendrite spines were visualized in stratum radiatum, focusing on multiple oblique secondary branches. Distal apical dendrite spines were visualized in stratum lacunosum moleculare, in multiple branch segments beyond the bifurcation of the primary apical dendrite and located greater than 350*μ*m from the soma. Thicker branches near the border of stratum radiatum and stratum lacunosum moleculare (Megias et al., 2001), terminal ends of the arbor, and branch points were avoided. Spines were counted posthoc using ImageJ by another individual blinded to genotype and PN identity. Spine density was derived from the spine number divided by segment length.

### Immunohistochemistry

Mice were anesthetized and transcardially perfused with 1X phosphate-buffered saline (PBS) followed by 4% paraformaldehyde (PFA). After decapitation, brains were removed from the skull and post-fixed in 4% PFA for 24 h. After rinsing in 1X PBS, 401m coronal slices were sectioned on a Leica VT1000s vibratome. Immunohistochemistry (IHC) was subsequently performed to detect amyloid (6E10, BioLegend #803004), phosphorylated tau (AT8, ThermoFisher Scientific #MN1020), hAPP (anti-Y188, Abcam #32136), and tau (HT7, ThermoFisher Scientific #MN1000).

For 6E10 and AT8 IHC, free-floating sections were permeabilized by rinsing thrice for 10 min in 1x PBS + 0.1% Triton, followed by incubation in blocking solution (PBS + 5% normal donkey serum) for 1 h at room temperature. Primary antibody incubation was carried out in blocking solution overnight at 4°C with 1:500 (6E10) or 1:100 (AT8) antibody dilution, followed by three washes in 1x PBS + 0.1% Triton. Secondary antibody incubation was performed in blocking solution for 2 h at room temperature (goat anti-mouse IgG1, Alexa Fluor 488, Life Technologies A21121; 1:500 dilution). Counterstaining was performed by adding NeuroTrace 640/660 (ThermoFisher Scientific #N21483) fluorescent Nissl stain at 1:200 dilution during secondary antibody incubations. Slices were rinsed thrice for 10 min in 1xPBS prior to mounting.

For anti-APP (Y188) IHC, slices were rinsed in 1X PBS three times for 5 min. Sections were blocked for 1 h at room temperature (RT) in 0.2% Triton X-100 and 5% normal goat serum (NGS). Slices were washed twice for 5 min in 1X PBS, followed by overnight incubation at 4°C in 3% NGS, 0.2% Triton X-100 and anti-Y188 (1:1000). Sections were rinsed thrice for 10 min in 1X PBS, immersed in 0.2% Triton X-100 and goat anti-rabbit AF 488 (1:500; Invitrogen #A11008) for 3h at RT, and rinsed thrice for 10 min in 1X PBS.

For HT7 IHC, sections were rinsed thrice for 5 min in 1X PBS and then immersed in 88% formic acid for 5 min, followed by a ddH20 rinse for 5-10 min. Sections were then rinsed twice for 5 min in 1X PBS, then immersed 1 h in blocking solution consisting of 5% NGS and 0.4% Triton X-100. Sections were stored overnight at 4°C in HT7 (1:1000). Sections were rinsed thrice for 15 min in 1X PBS and then immersed in goat anti-mouse AF 594 antibody solution (1:1000; Invitrogen #A11032) for 3 h at RT. Lastly, sections were rinsed thrice for 15 min in 1X PBS.

For Y188 and HT7 labelings, sections were coverslipped with Fluoromount-G mounting medium containing DAPI (Invitrogen #00-4959-52) and viewed with an upright Zeiss LSM 800 Axio Imager confocal microscope (Carl Zeiss Microscopy, LLC) and associated Zen Blue 2.3 software. For 6E10 and AT8 labelings, images were acquired on a Zeiss LSM 700 laser scanning confocal microscope with Zen 2012 SP5 FP3 black edition software, using either a Zeiss Fluar 5x/0.25 objective (0.5 zoom, pixel size: 2.5 x 2.5 μm2) or a Zeiss Plan-Apochromat 20X/0.8 objective (1.0 zoom, pixel size: 0.3126 x 0.3126 μm2).

### Data Analysis

Electrophysiological measurements were quantified using Clampfit. Statistical analysis was performed with Prism (Graphpad). Statistical errors are shown as standard errors of the mean. For comparing singular means, 2-way ANOVA was used to evaluate differences as a function of age/genotype or cell type. P-values were adjusted post-hoc for multiple comparisons with Bonferroni’s correction. Comparison between curves was also accomplished via 2-way ANOVA, with p-value relating to the hypothesis that age/genotype (or cell type) drove a difference between the overall curves.

## ACKNOWLEDGEMENTS

Grant support for this work was from NIH RF1AG072507, R01AG072507 (A.V.M.), R21AG070880 (A.V.M.), T32MH020004 (A.V.M.), Alzheimer’s Association Clinical Fellowship (A.V.M.), Blas Frangione Foundation (A.V.M.), Leon Levy Foundation (A.V.M.), Fondazione Cariplo (B.S.). We thank Steven Siegelbaum for his valuable input and guidance.

## FIGURE LEGENDS

**Supplemental Figure 1.**
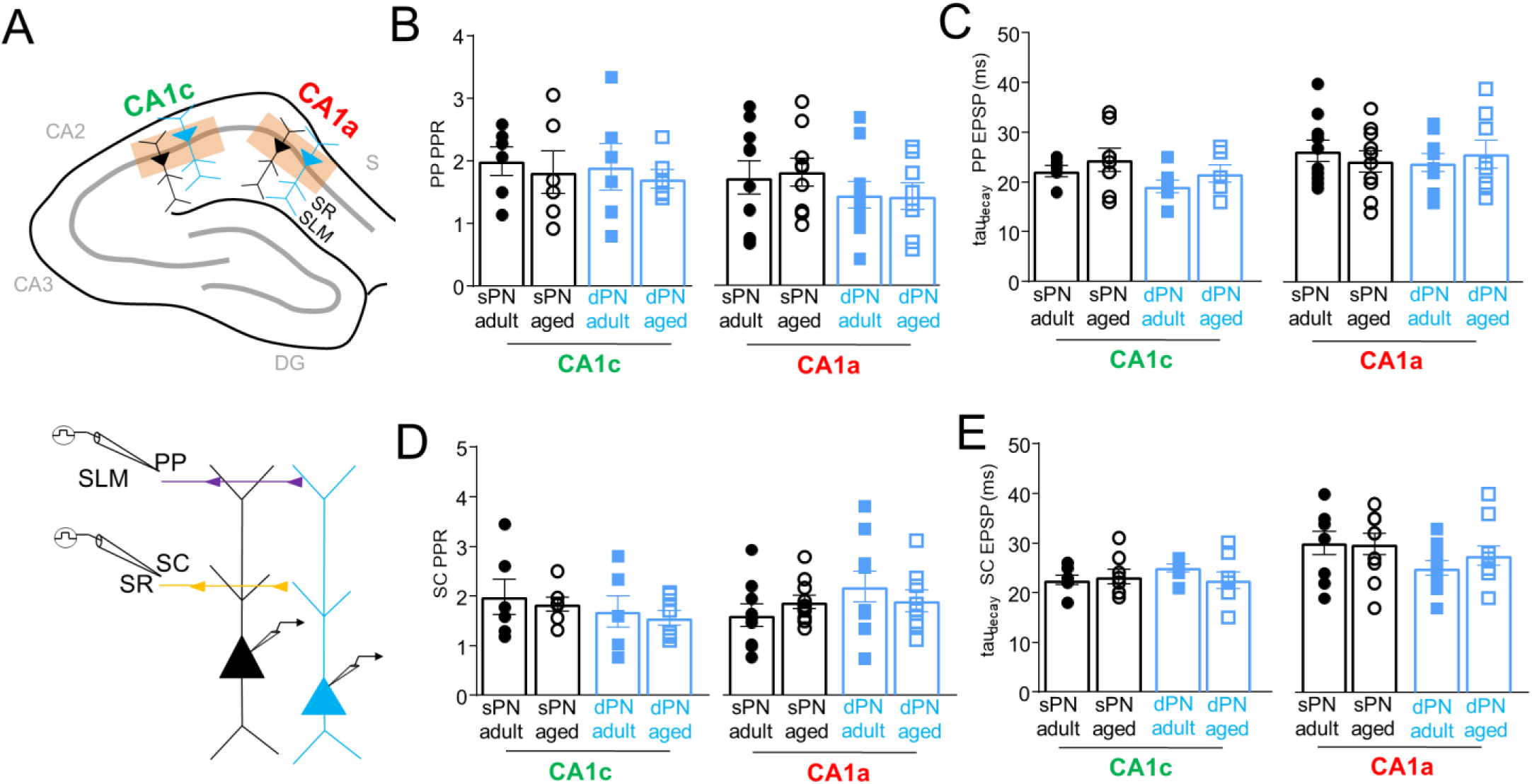
Impact of aging on paired pulse ratio and EPSP decay time constant of PP and SC inputs. (A) In adult and aged male mice, whole cell recordings were targeted to (above) sPNs and dPNs in CA1c and CA1a, with (below) PP and SC responses elicited by extracellular stimulation of SLM and SR, respectively. The (B) paired pulse ratio (PPR) and (C) decay time constant (tau_decay_) of the PP EPSP were measured in sPNs/dPNs of CA1c (aged: n = 6-8 from 5-6 mice/group; adult: n = 6-7 from 3-4 mice/group) and CA1a (aged: n = 8-10 from 5-6 mice/group; adult: n = 9-10 from 6 mice/group). The (D) PPR and (E) tau_decay_ of the SC EPSP were measured in sPNs/dPNs of CA1c (aged: n = 7-8 from 5-6 mice/group; adult: n = 6-7 from 3-4 mice/group) and CA1a (aged: n = 8-10 from 5-7 mice/group; adult: n = 9-10 from 5-6 mice/group).

**Supplemental Figure 2.**
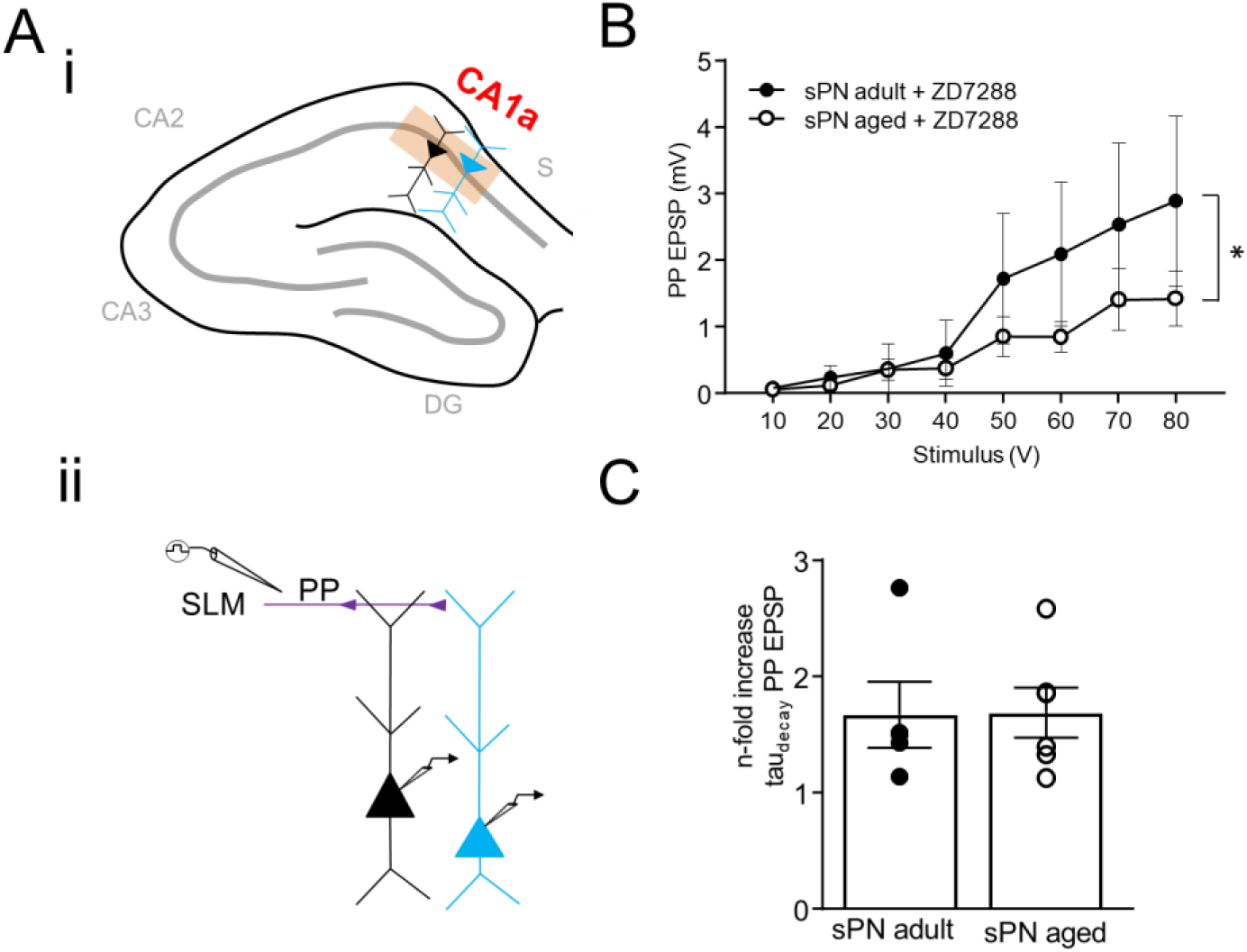
Impact of HCN current blockade on PP EPSPs in CA1a sPNs of adult and aged mice. (A) In adult and aged male mice, whole cell recordings were targeted (i) sPNs in CA1a, with (ii) PP responses elicited by extracellular stimulation of SLM. (B) In the setting of 10*μ*m ZD7288, the PP EPSP input-output curve in adult and aged mice (n = 5-6 from 3 mice/group) (C) n-fold change in tau_decay_ of PP EPSP with addition of 10*μ*m ZD7288 in adult and age mice (n = 5-6 from 3 mice/group). For all data above, error bars are +/-SE. *p < 0.05

**Supplemental Figure 3.**
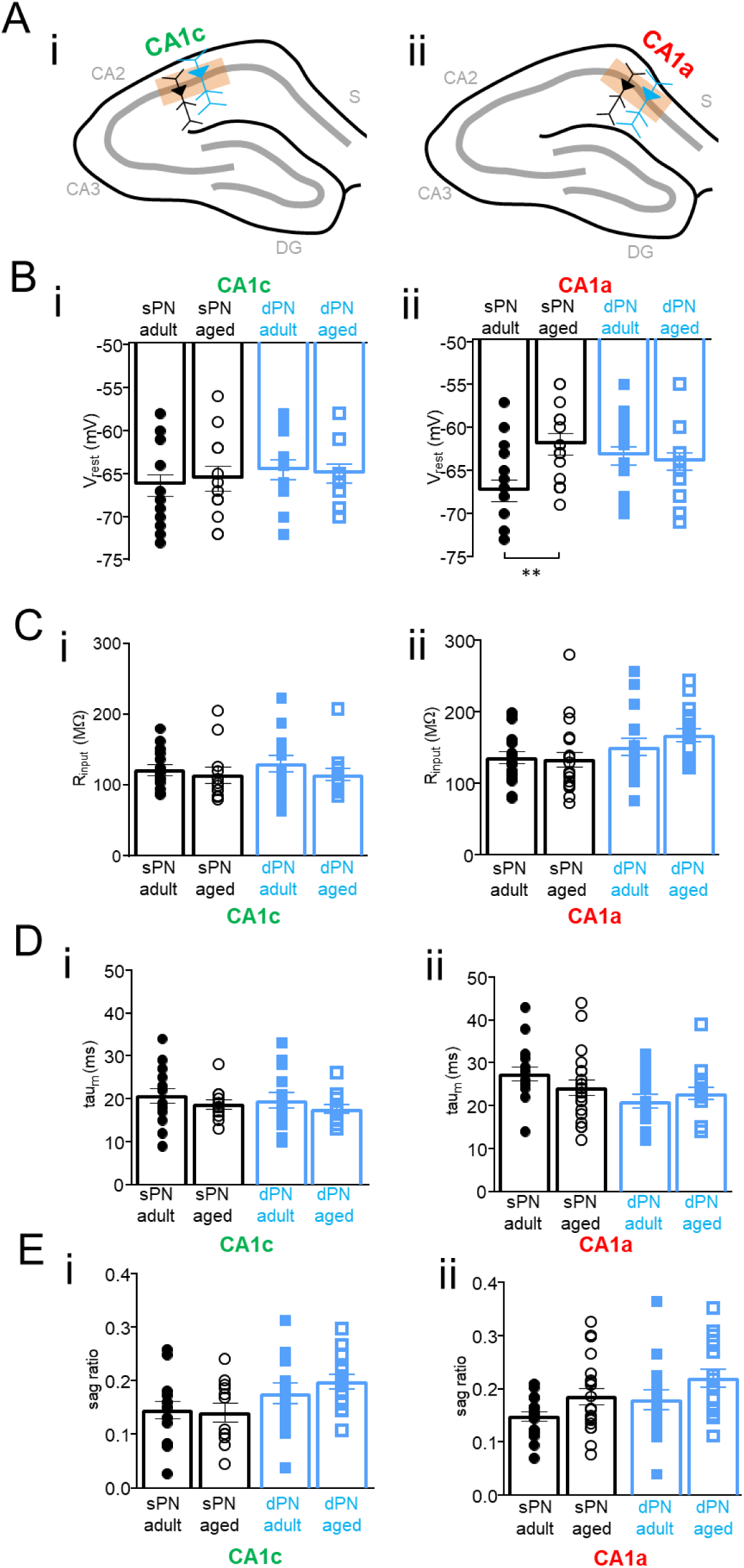
Aging-related changes in subthreshold intrinsic properties. (A) Whole cell recordings were targeted to sPNs and dPNs in (i) CA1c (aged: n = 12-14 from 8 mice/group; adult: n = 14-15 from 8-9 mice/group) and (ii) CA1a (aged: n = 17-20 from 6-7 mice/group; adult: n = 17 from 10 mice/group) of adult and aged male mice to measure resting membrane potential V_rest_ and other intrinsic properties. (B) Resting membrane potential V_rest_ in (i) CA1c and (ii) Ca1a. (C) R_input_ in (i) Ca1c and (ii) CA1a, (D) tau_m_ in (i) Ca1c and (ii) CA1a, and (E) sag ratio in (i) Ca1c and (ii) CA1a. For all data above, error bars are +/-SE. **p < 0.01.

**Supplemental Figure 4.**
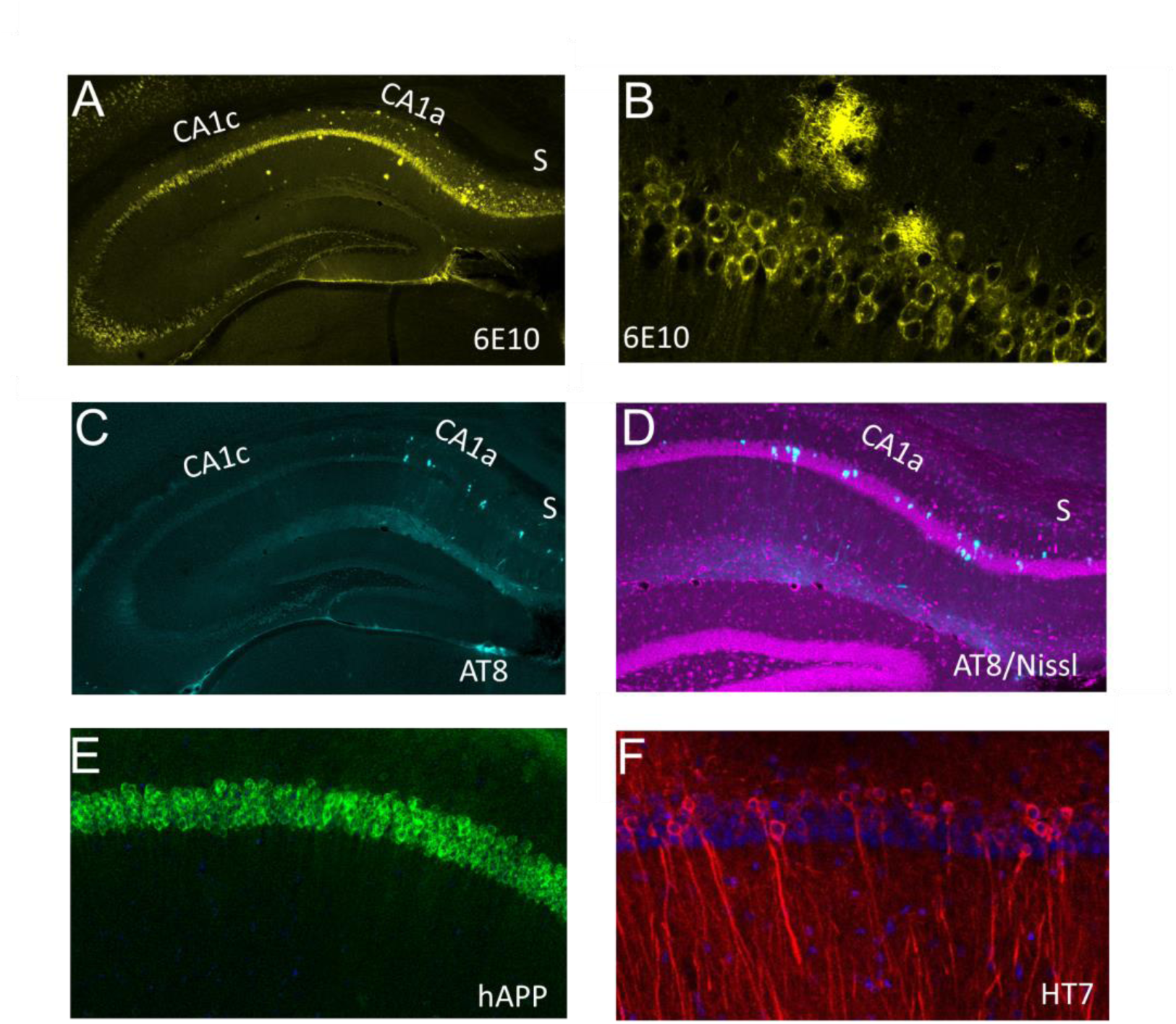
Immunohistochemical characterization of aged 3xTg-AD mice. (A) 6E10 staining shows extracellular amyloid plaque primarily in CA1a. (B) Magnification of 6E10 staining in CA1a. (C) AT8 staining shows intracellular phosphorylated tau pathology primarily in CA1, with (D) closeup of CA1a with Nissl counterstain revealing AT8 staining in superficial and deep PN layers. (E) hAPP staining in CA1a shows uniform APP transgene expression in superficial and deep PN layers. (F) HT7 staining in CA1a shows similar tau transgene expression in superficial and deep PN layers.

**Supplemental Figure 5.**
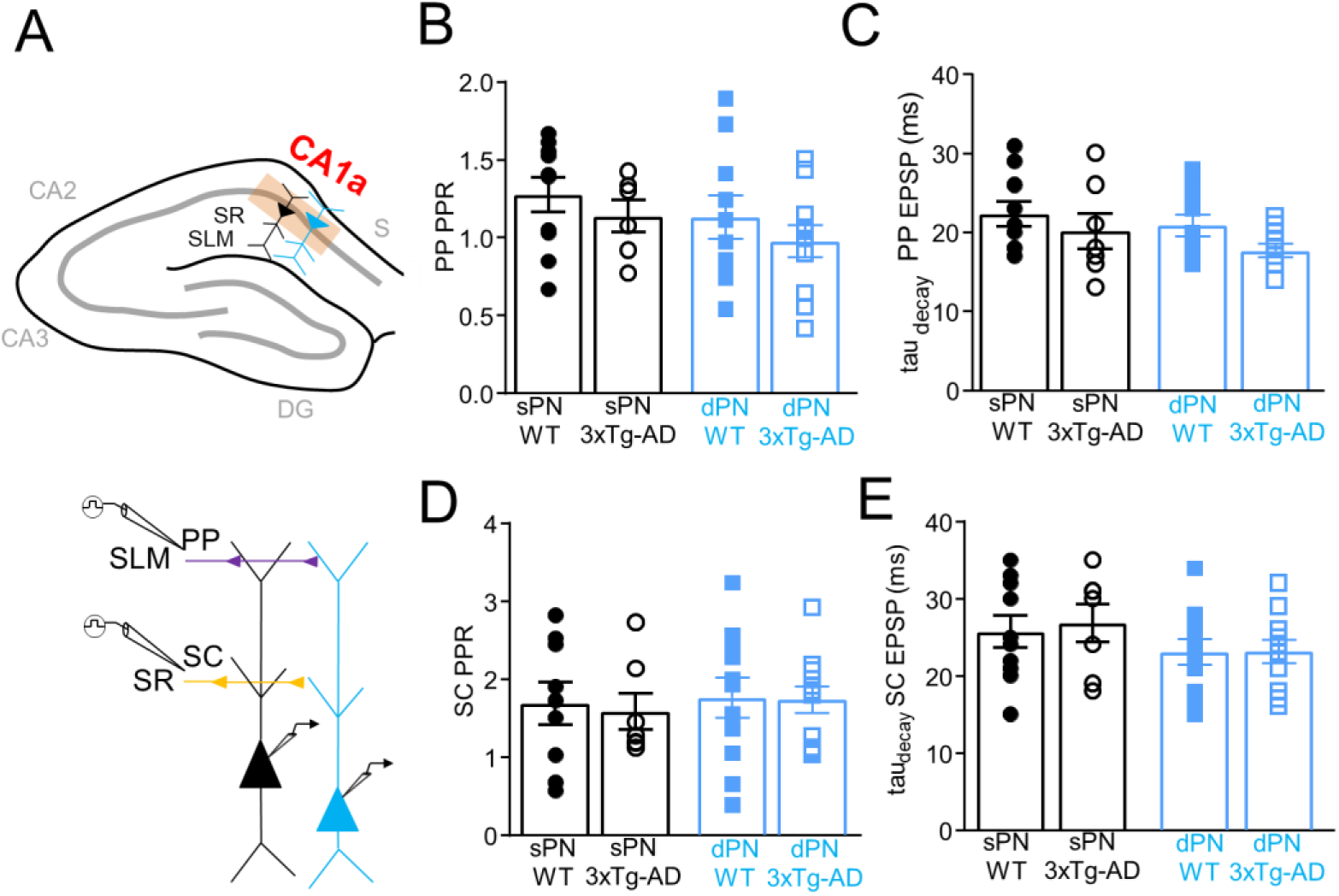
Paired pulse ratio and EPSP decay time constant of PP and SC inputs in aged 3xTg-AD and WT mice. (A) In aged female 3xTg-AD and WT mice, whole cell recordings were targeted to (above) sPNs and dPNs in CA1a, with (below) PP and SC responses elicited by extracellular stimulation of SLM and SR, respectively. (B) PPR and (C) tau_decay_ of the PP EPSP (AD: n = 6-11 from 5 mice/group; WT: n = 10-11 from 5 mice/group). (D) PPR and (E) tau_decay_ of the SC EPSP (AD: n = 7-11 from 5 mice/group; WT: n = 10-11 from 5 mice/group).

**Supplemental Figure 6.**
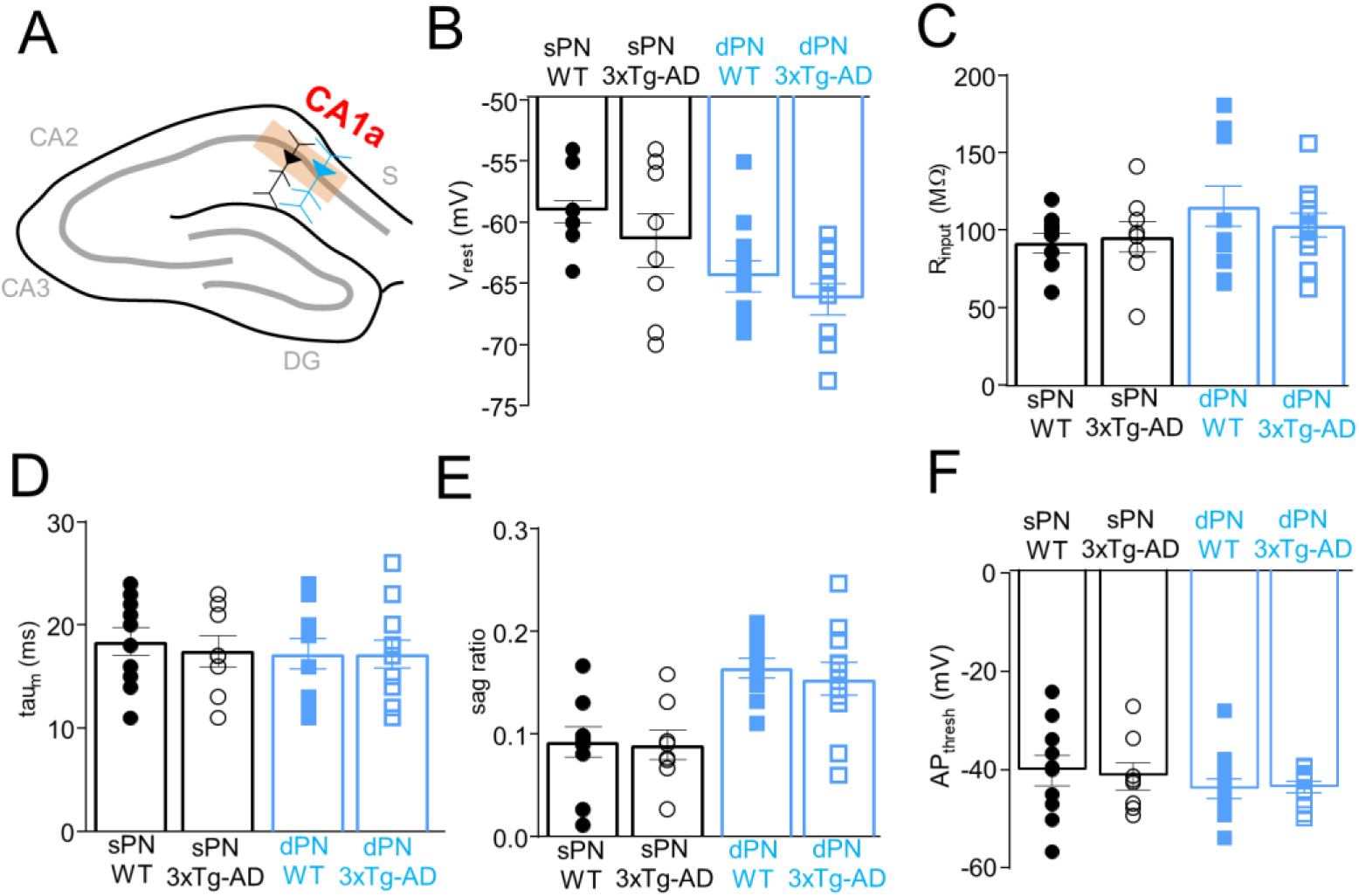
Changes in intrinsic properties of CA1a PNs in aged 3xTg-AD versus WT mice. (A) Whole cell recordings were targeted to CA1a sPNs and dPNs of aged 3xTg-AD and WT mice to measure subthreshold and suprathreshold intrinsic properties, including (B) V_rest_, (C) R_input_, (D) tau_m_, (E) sag ratio, (F) AP threshold. For all data above, n = 8-11/group from 5 mice/group. Error bars are +/-SE.

